# Reforestation without Belowground Recovery: Sediment Microbiomes and Organic Carbon in Caribbean Mangroves

**DOI:** 10.64898/2026.10.03.756322

**Authors:** Hansel B. Lopez-Chevalier, Andy N. Avelino-Perez, Juleidy E. Suriel-Jaime, Leslys Garcia-Encarnacion, Julian A. Cabrera-Hernandez

**Author notes:** Correspondence: Hansel B. Lopez-Chevalier.

## Abstract

Mangrove reforestation is expanding across the Caribbean, but restoration success is still judged almost exclusively by tree survival and canopy cover. Whether the sediment microbiome, which governs carbon burial and nutrient cycling, follows the recovery of the vegetation remains largely unknown. We combined 16S rRNA gene metabarcoding (V4, Illumina MiSeq) with CHNS elemental analysis, pH and salinity measurements to compare sediment prokaryotic communities along 0–100 cm profiles in a reference, a reforested (*Rhizophora mangle* plantings since 2019) and an altered mangrove within the Las Dunas de Las Calderas Natural Monument, Dominican Republic (39 cores pooled into 28 depth-resolved composite samples). Reference sediments contained 2.5–5.5-fold more organic carbon (median 7.5% vs 3.0% in altered and 1.3% in reforested sediments), had higher C/N ratios and were less alkaline (pH 8.2 vs 8.8). Reference communities were more diverse and even, and differed in composition from both disturbed stands (pairwise PERMANOVA, R^2^ ≈ 0.09, adjusted *P* = 0.0015), whereas reforested and altered communities were statistically indistinguishable (R^2^ = 0.04, *P* = 0.25). Disturbed sediments were dominated by Cyanobacteriota (≈41–45% of reads), which tracked high pH, whereas reference sediments were enriched in Chloroflexota, Thermodesulfobacteriota and Thermoproteota, lineages linked to anaerobic degradation of complex organic matter and sulfate reduction. Depth structured communities by separating surface from subsurface layers. Sediment organic carbon and nitrogen were the only significant constraints in distance-based redundancy analysis. These results indicate that, within a few years of the first plantings (2019), reforestation had not restored the carbon-rich, anaerobic sediment habitat of the reference forest. We propose sediment organic carbon and microbial community structure as complementary indicators for mangrove restoration monitoring in the Caribbean.

## Introduction

Mangroves are among the most carbon-dense ecosystems on Earth. Although they occupy a small fraction of the world’s coastlines, most of their carbon is stored below-ground, in waterlogged sediments where oxygen limitation slows the decomposition of root-derived and lignin-rich organic matter^1–4^. Mangroves also stabilise shorelines, retain nutrients and sustain coastal fisheries^5,6^. Despite this value, they continue to be lost to coastal development, aquaculture, logging and hydrological alteration^5,6^. In the Dominican Republic, mangrove cover declined from about 25,200 to 18,400 ha between 1969 and 2012^7^, and mangroves have been declared of high national interest for protection and rehabilitation since 1987^8^.

Planting propagules or seedlings has become the default restoration strategy, but its outcomes are uneven. Many projects achieve canopy establishment while failing to restore hydrology, sediment properties or ecosystem functions^9–11^. A recurring problem is the use of single-species plantings outside the species’ hydrological and salinity niche, which is particularly common with *Rhizophora mangle* in the Caribbean^12,13^. Because restoration success is still evaluated mainly through aboveground metrics, it remains unclear whether planted stands recover the belowground processes that underpin carbon storage^11,14,15^.

Sediment microorganisms control those processes. Bacteria and archaea mediate organic-matter hydrolysis and fermentation, sulfate reduction, methanogenesis and methane oxidation, nitrogen fixation and denitrification, ultimately determining whether buried carbon is preserved or respired^16–19^. Mangrove sediments host distinctive communities rich in anaerobic lineages such as sulfate-reducing Thermodesulfobacteriota/Desulfobacterota, Chloroflexota and diverse archaea, vertically organised along redox gradients^18,20–22^. Because these communities respond to organic carbon, nutrients, pH, salinity and oxygen, they integrate the biogeochemical state of the sediment and have been proposed as indicators of mangrove conservation status^14,23^.

Evidence on microbial recovery after mangrove restoration is mixed. Degradation consistently reduces microbial diversity and the abundance of taxa involved in carbon, nitrogen and sulfur cycling^24–26^. Some restored stands converge towards the composition and multifunctionality of conserved forests^27–29^, whereas others remain distinct even after years to decades^15,30,31^. Most of this evidence comes from Asia, Brazil and Mexico. In the insular Caribbean, and in arid mangrove settings where carbon stocks and microbial communities may behave differently^32,33^, sediment microbiomes of restored mangroves have not been studied.

Las Calderas Bay, on the arid south coast of the Dominican Republic, offers a natural experiment. Within the Las Dunas de Las Calderas Natural Monument, a relatively intact mangrove persists next to stands degraded by clearing and burning^34^ and to areas reforested with *R. mangle* since 2019^35^, all sharing the same climate, tidal regime and parent material. Here we characterised sediment prokaryotic communities and physicochemistry across four depth intervals (0–100 cm) in reference, reforested and altered stands (Fig. 1). We asked (i) whether sediment properties and microbial diversity and composition of the reforested stand converge towards the reference or remain similar to the altered stand, (ii) how depth structures these communities, and (iii) which environmental variables best explain community variation. We hypothesised that, if reforestation had restored sediment function, the reforested microbiome would resemble the reference community and diverge from the altered one.

**Figure 1:**
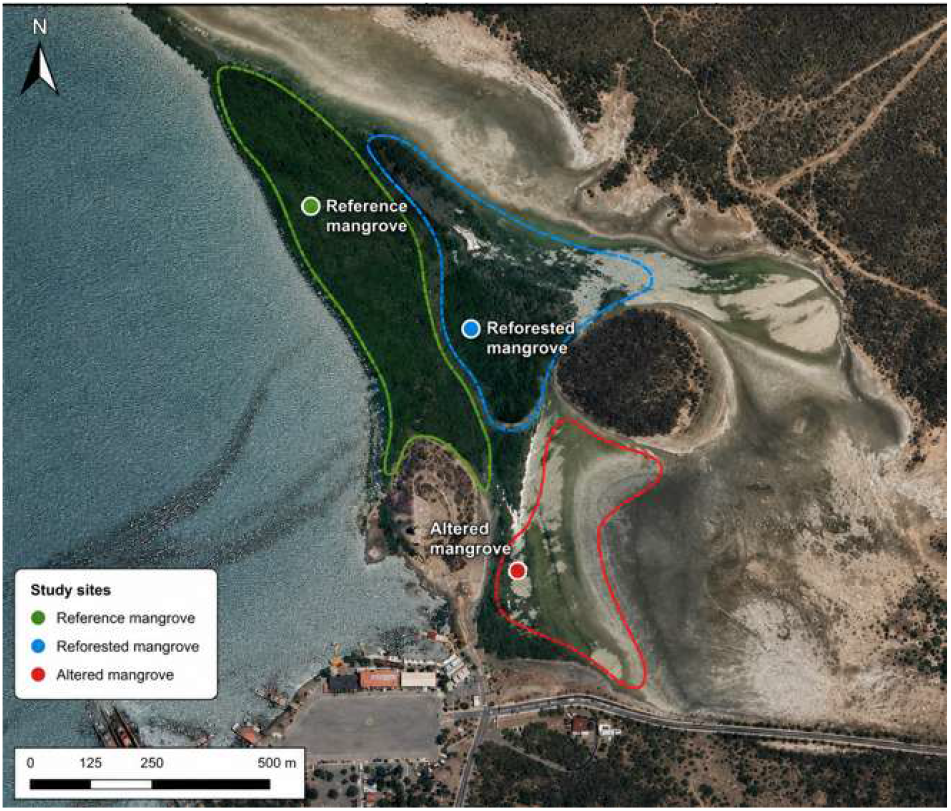
Study area in Las Calderas Bay, Peravia province, Dominican Republic. Reference (green), reforested (blue) and altered (red) mangrove stands within the Las Dunas de Las Calderas Natural Monument. Outlines delimit each stand and circles indicate the sampling areas. Basemap: Google Earth satellite imagery, July 2026. Scale bar, 500 m.

## Results

### Sequencing controls

All 28 sediment libraries were rarefied to 14,536 reads per sample. The ZymoBIOMICS D6300 mock community was recovered in proportions closely matching its theoretical 16S rRNA gene composition, with all eight bacterial members detected and deviations of ≤0.7 percentage points from the expected abundances (Supplementary Fig. S1; Supplementary Table S5). The negative control yielded a low, diffuse signal of environmental phyla, mainly Pseudomonadota, Verrucomicrobiota and Patescibacteria, and contained none of the mock taxa. Contaminant ASVs identified from this control were removed before analysis^36^.

### Reference sediments are carbon-rich and less alkaline

Sediment properties separated the reference stand from both disturbed stands (Fig. 2; Supplementary Tables S1, S3). Total organic carbon (TOC) was highest in the reference stand (median 7.5%, range 5.2–9.3%), intermediate in the altered stand (3.0%, 2.2–4.6%) and lowest in the reforested stand (1.3%, 1.1–2.1%). All pairwise contrasts were significant (Wilcoxon rank-sum, BH-adjusted *P* ≤ 0.004). The C/N ratio followed the same order (median 9.9, 3.2 and 1.7; adjusted *P* ≤ 0.004). Reference sediments were less alkaline (pH 8.13–8.44) than reforested (8.51–9.01) and altered (8.50–9.10) sediments (adjusted *P* = 0.010 for both), whereas the two disturbed stands did not differ in pH. Nitrogen was highest in the altered stand (median 0.95%) and significantly higher there than in the reforested stand (0.82%; adjusted *P* = 0.020). Sulfur was below detection in 9 of 12 altered samples and was lower there than in reforested (adjusted *P* = 0.004) and reference sediments (adjusted *P* = 0.020). Hydrogen and salinity (NaCl 2.1–8.5%) did not differ among stands. Inorganic carbon followed the same order as TOC (median 1.32%, 0.60% and 0.30% in reference, altered and reforested sediments) but represented a similar fraction of total carbon in all three stands (median 17–18%; range 10–25%), so the differences in carbon among stands were driven by organic carbon (Supplementary Table S1).

**Figure 2:**
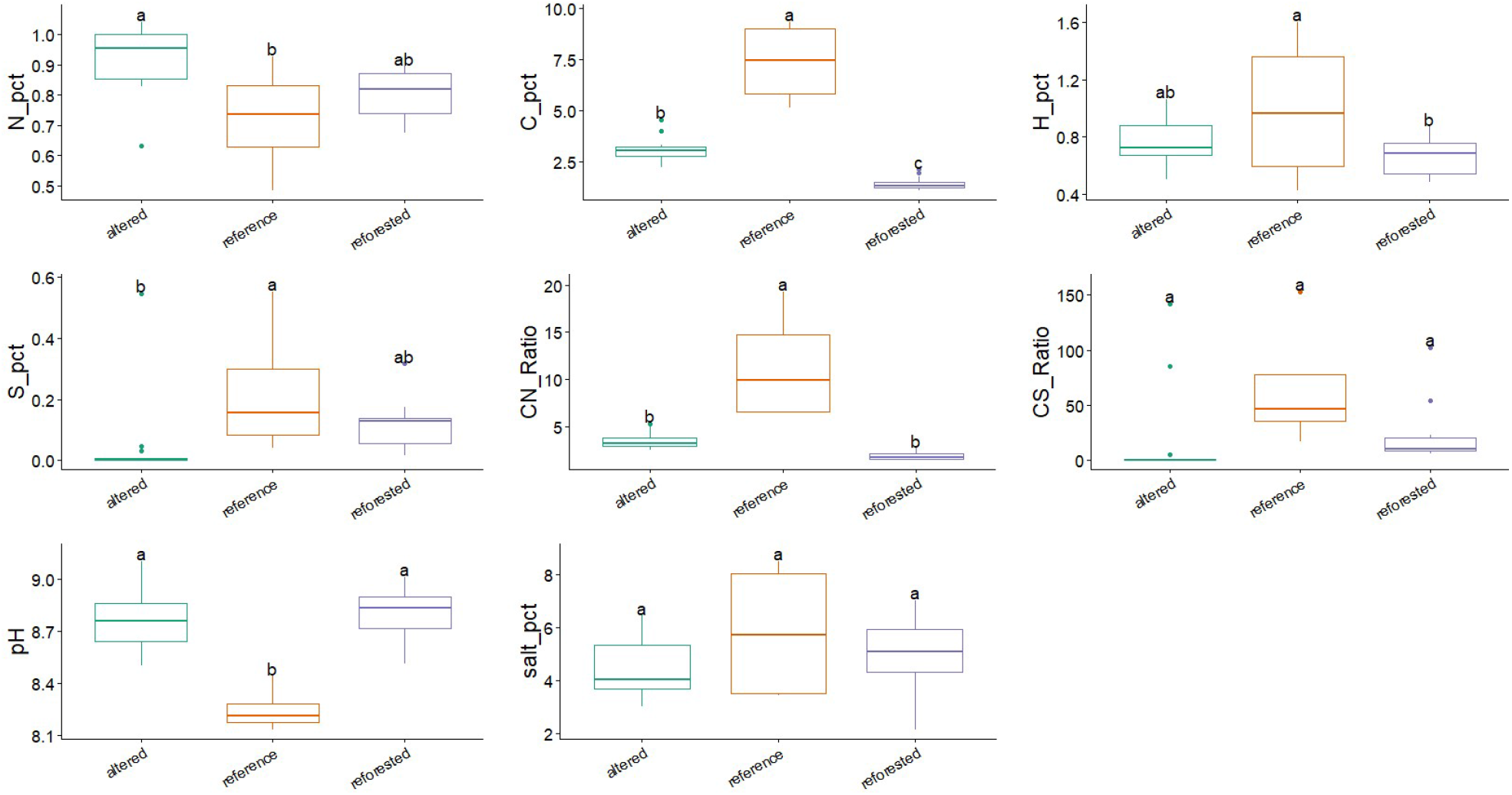
Sediment physicochemistry of the three mangrove stands. Nitrogen (N), total organic carbon (C), hydrogen (H) and sulfur (S) contents (% dry weight), C/N and C/S ratios, pH and salinity (NaCl, %) in altered (n = 12), reference (n = 4) and reforested (n = 12) composite samples pooled across depths. Boxes show the median and interquartile range; whiskers extend to 1.5 × IQR. Different letters indicate significant differences (ANOVA followed by Tukey’s HSD, *P* < 0.05). Pairwise Wilcoxon rank-sum tests are given in Supplementary Table S3.

Depth profiles also differed (Supplementary Figs. S4, S5). In the reference stand, TOC increased from 6.0% at 0– 15 cm to 9.3% at 50–100 cm while nitrogen decreased from 0.93% to 0.48%, so that C/N rose from 6.5 to 19.3 with depth. Sulfur also increased with depth in the reference stand (0.04% to 0.55%). By contrast, TOC was nearly constant with depth in the reforested (1.1–2.1%) and altered (2.2–4.6%) stands. Salinity increased with depth in all three stands.

### Cyanobacteriota dominate the disturbed sediments

Communities were dominated by Cyanobacteriota, Acidobacteriota, Chloroflexota and Pseudomonadota, followed by Thermodesulfobacteriota, Actinomycetota, Halobacteriota, Thermoproteota and Methanobacteriota (Fig. 3). Reforested and altered sediments had strikingly similar profiles: Cyanobacteriota accounted for ≈44% and ≈41% of reads, followed by Acidobacteriota (≈17% and ≈26%). In the reference stand, Cyanobacteriota represented only ≈14% of reads, and the community was more evenly distributed among Acidobacteriota (≈19%), Chloroflexota (≈16%), Thermodesulfobacteriota (≈10%) and Pseudomonadota (≈10%) (Fig. 3A).

**Figure 3:**
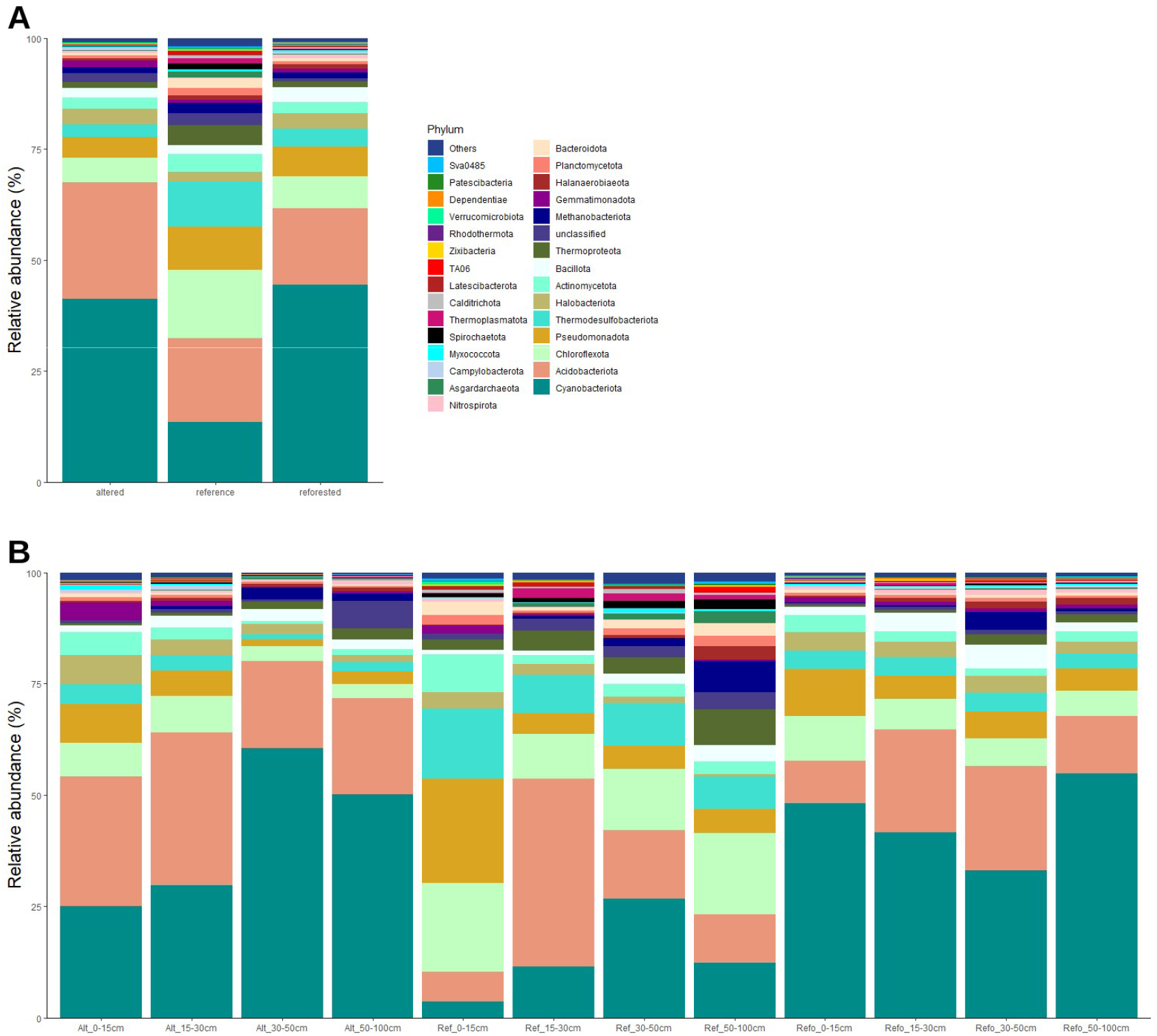
Phylum-level composition of sediment prokaryotic communities. **(A)** Mean relative abundance of phyla in altered, reference and reforested stands. **(B)** Mean relative abundance by stand and depth interval (Alt, altered; Ref, reference; Refo, reforested). Data were rarefied to 14,536 reads per sample.

Cyanobacteriota were significantly depleted in the reference stand relative to both disturbed stands, whereas Chloroflexota, Thermodesulfobacteriota and Thermoproteota were significantly enriched (Wilcoxon rank-sum, *P* < 0.05 to *P* < 0.01; Fig. 6A). Cyanobacteriota remained abundant down to 50–100 cm in the reforested (≈55%) and altered (≈50%) profiles, peaking at 30–50 cm in the altered stand (≈61%) (Fig. 3B). The deepest reference layer was distinguished by a larger contribution of archaeal lineages (Methanobacteriota, Thermoproteota, Asgardarchaeota) and Halanaerobiaeota. Pseudomonadota and Acidobacteriota did not differ significantly among stands or depths, and a parametric comparison of the four dominant phyla detected only the higher abundance of Chloroflexota in the reference stand (Supplementary Fig. S2).

### Lower diversity and evenness outside the reference stand

Shannon diversity and Pielou’s evenness were significantly higher in the reference stand than in reforested and altered stands (ANOVA, Tukey’s HSD *P* < 0.05; median Shannon ≈5.6 vs ≈4.6 and ≈4.1; median evenness ≈0.85 vs ≈0.71 and ≈0.68). Observed richness and Chao1 did not differ (Fig. 4), and the close agreement between the two indices indicates that sequencing depth captured most of the richness present^37^. Diversity loss in the disturbed stands therefore reflected greater dominance by a few taxa rather than loss of taxa. Reforested and altered stands did not differ in any index. Diversity was most variable in the altered stand, where the 30–50 cm layer had the lowest richness of the dataset (median ≈130 observed ASVs) and the 50–100 cm layer the lowest evenness (Supplementary Fig. S3).

**Figure 4:**
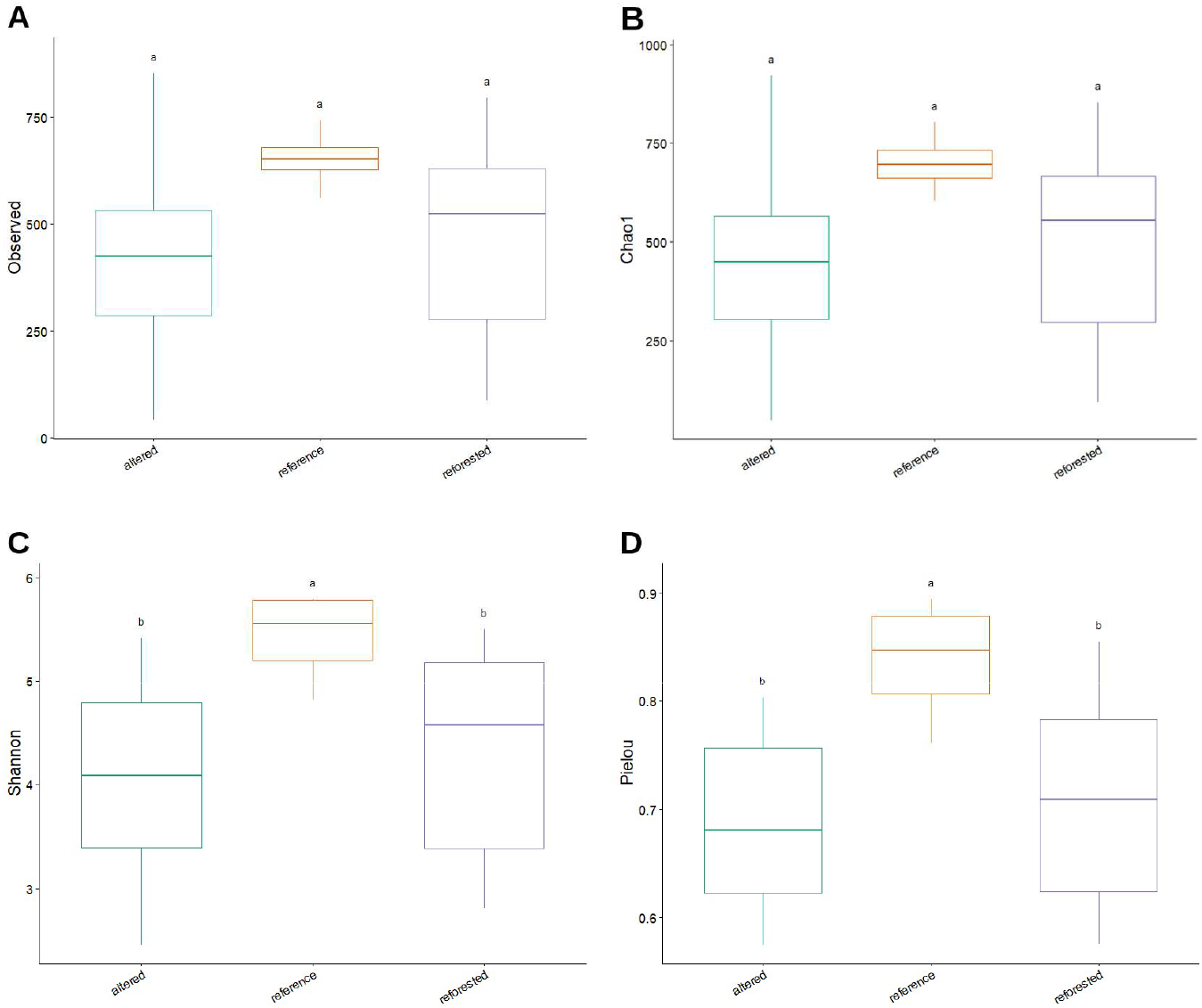
Alpha diversity of sediment prokaryotic communities. **(A)** Observed ASV richness, **(B)** Chao1, **(C)** Shannon index and **(D)** Pielou’s evenness in altered (n = 12), reference (n = 4) and reforested (n = 12) sediments. Different letters denote significant differences (ANOVA and Tukey’s HSD, *P* < 0.05).

### Reforested and altered communities are indistinguishable

NMDS of Bray–Curtis dissimilarities (stress = 0.14) placed the four reference samples in a compact cluster, whereas reforested and altered samples overlapped broadly and were more dispersed (Fig. 5A). Pairwise PERMANOVA confirmed that the reference community differed from both the reforested (*F* = 1.38, R^2^ = 0.090, adjusted *P* = 0.0015) and the altered community (*F* = 1.43, R^2^ = 0.093, adjusted *P* = 0.0015). Reforested and altered communities did not differ (*F* = 1.02, R^2^ = 0.044, *P* = 0.252) (Supplementary Table S2).

**Figure 5:**
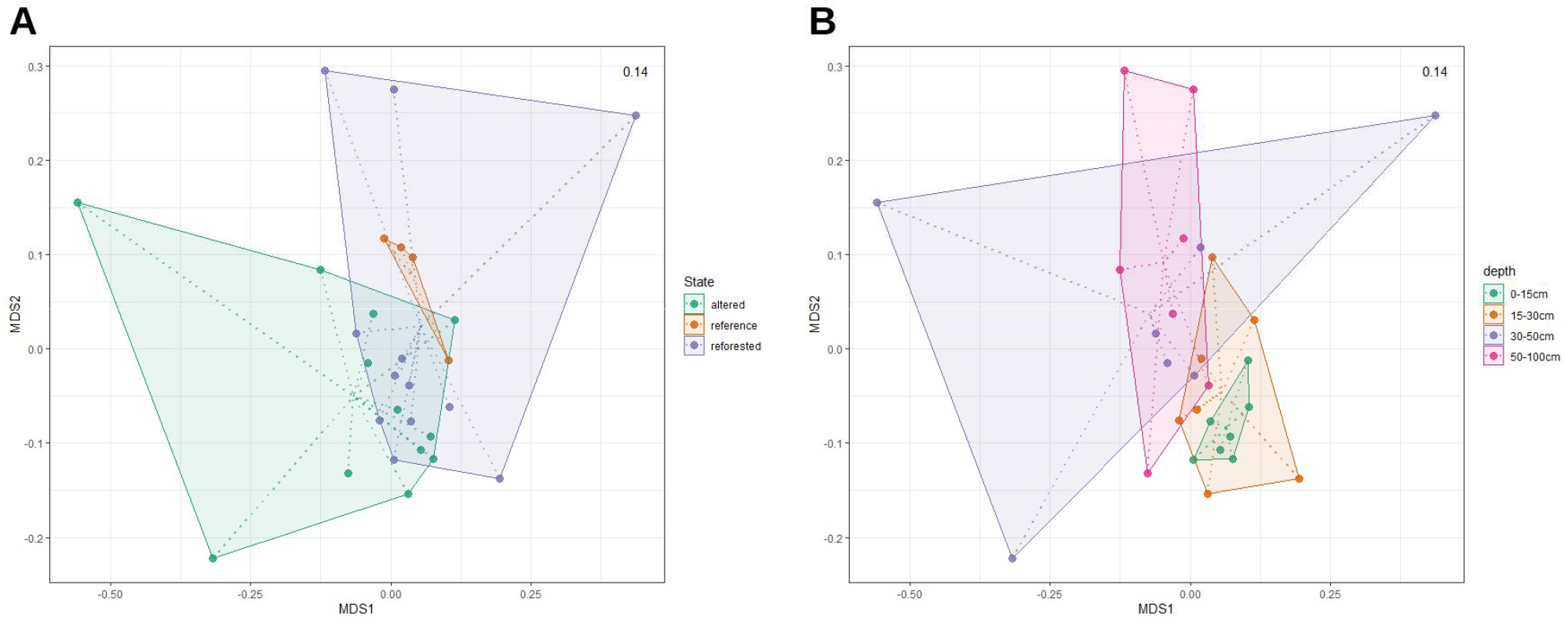
Community structure by stand and depth. Non-metric multidimensional scaling (NMDS) of Bray–Curtis dissimilarities among the 28 composite samples (stress = 0.14), coloured by **(A)** stand and **(B)** depth interval. Polygons are convex hulls; dotted lines connect samples to their group centroid.

**Figure 6:**
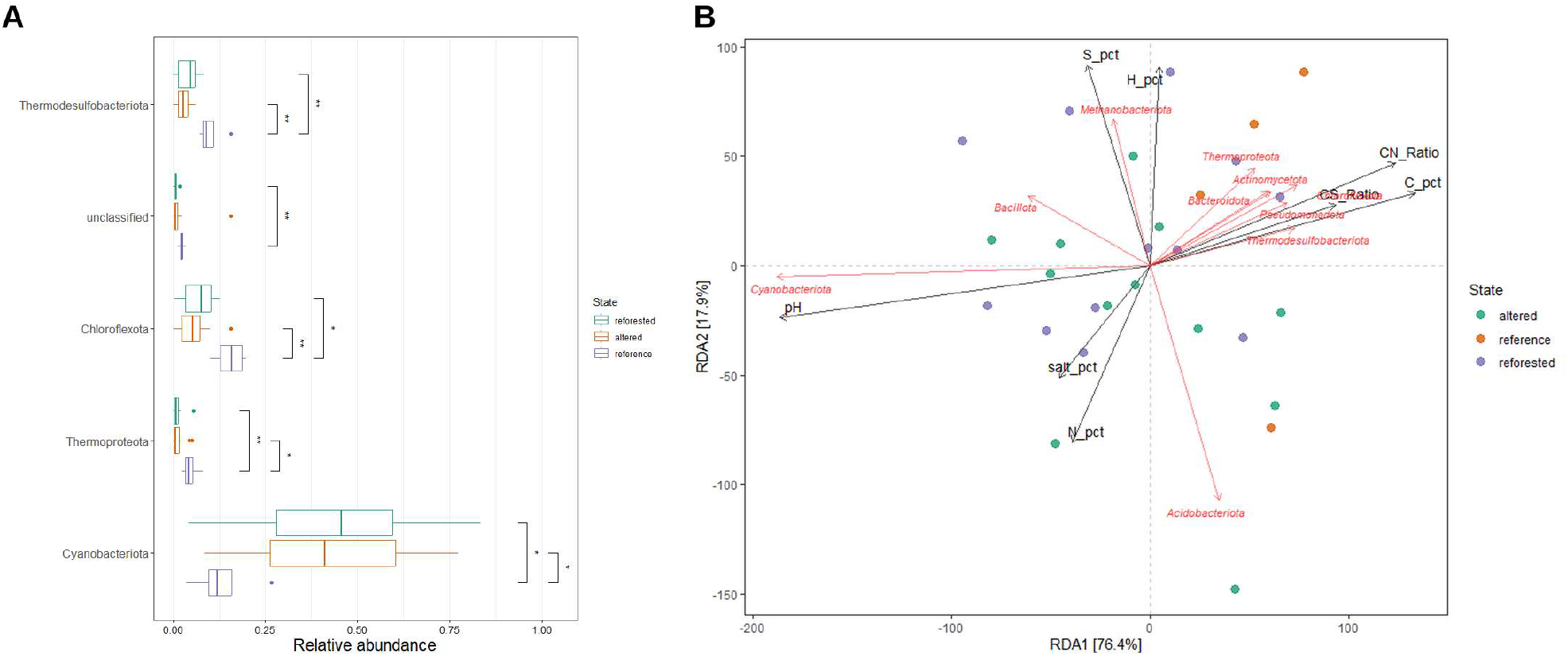
Differentially abundant phyla and their environmental associations. **(A)** Phyla whose relative abundance differed significantly among stands (pairwise Wilcoxon rank-sum tests; ^*^ *P* < 0.05, ^**^ *P* < 0.01). “Unclassified” denotes ASVs without phylum assignment. **(B)** Redundancy analysis of phylum-level relative abundances constrained by sediment variables. Black arrows, environmental variables; red arrows, phyla.

Depth structured the communities mainly by separating the surface layer from deeper sediments (Fig. 5B). Surface (0–15 cm) communities differed from those at 30–50 cm (R^2^ = 0.091, adjusted *P* = 0.021) and 50–100 cm (R^2^ = 0.091, adjusted *P* = 0.012), and marginally from 15–30 cm (unadjusted *P* = 0.037, adjusted *P* = 0.074). No differences were detected among the subsurface layers.

### Carbon and nitrogen are the main environmental drivers

Distance-based redundancy analysis (dbRDA) with eight sediment variables accounted for 29.9% of total Bray–Curtis inertia (Fig. 7). In sequential permutation tests, only organic carbon (*F* = 1.20, *P* = 0.005) and nitrogen (*F* = 1.09, *P* = 0.047) explained significant variation. H, S, C/N, C/S, pH and salinity did not (*P* ≥ 0.40; Supplementary Table S4). Only the first constrained axis was significant (*F* = 1.44, *P* = 0.008). dbRDA1 followed the organic carbon and C/N gradient and separated reference samples from most reforested and altered samples, whereas pH and nitrogen vectors pointed in the opposite direction.

**Figure 7:**
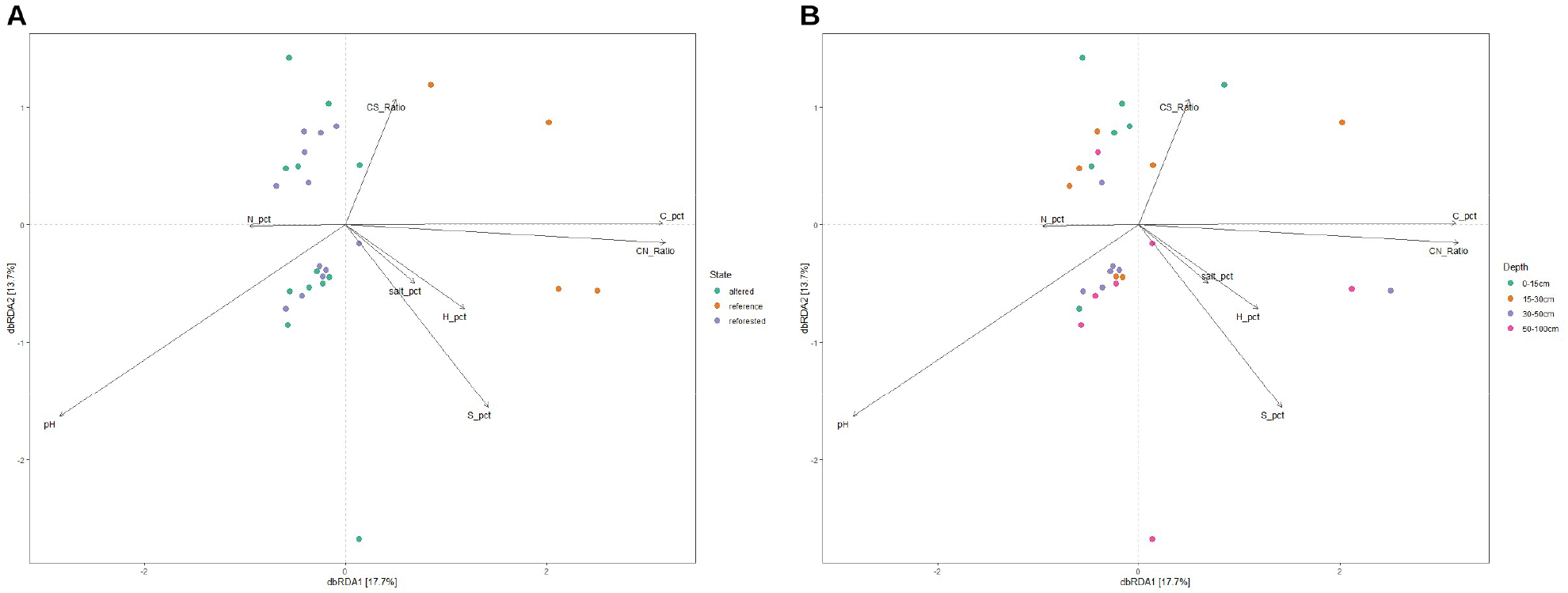
Distance-based redundancy analysis (dbRDA). dbRDA of Bray–Curtis dissimilarities constrained by N, total organic carbon (C), H and S contents, C/N and C/S ratios, pH and salinity (log_10_(x + 1)-transformed), with samples coloured by **(A)** stand and **(B)** depth interval. Axis labels give the percentage of constrained variance. Only C (*P* = 0.005) and N (*P* = 0.047) were significant in sequential permutation tests (999 permutations; Supplementary Table S4).

A phylum-level redundancy analysis showed the same structure (Fig. 6B). The first axis (76.4% of constrained variance) opposed Cyanobacteriota, aligned with pH, to Thermodesulfobacteriota, Chloroflexota, Pseudomonadota, Actinomycetota, Bacteroidota and Thermoproteota, aligned with carbon, C/N and C/S. Acidobacteriota were associated with nitrogen and salinity, and Methanobacteriota with sulfur and hydrogen. Phylum–environment correlations within each stand are shown in Supplementary Fig. S6.

## Discussion

### Reforestation without belowground recovery

A few years after the first plantings in 2019, the reforested stand of Las Calderas had not recovered either the sediment carbon or the prokaryotic community of the adjacent reference mangrove. On every axis we measured — carbon, C/N, pH, diversity, evenness, community composition and the abundance of anaerobic lineages — the reforested sediments resembled the altered stand rather than the reference forest, and the two disturbed communities were statistically indistinguishable. Our results therefore reject the hypothesis of convergence and agree with syntheses showing that planting alone does not restore ecosystem function while the drivers of degradation persist^9,11,15^.

This outcome differs from studies in which restored mangroves recovered much of the diversity and function of conserved forests^27–29^. Two features of our system may explain the contrast. First, the reforested stand is young. Microbial and carbon recovery after mangrove establishment can take one to several decades^30,31^, and sediment organic matter accumulates far more slowly than canopy cover. Second, reforestation in Las Calderas has relied on single-species *R. mangle* plantings^35^, a species whose survival and growth depend strongly on flooding, salinity and tidal position^12,13^. Where hydrology and sediment conditions are not restored, planted seedlings provide little of the root biomass and litter that feed the anaerobic food web of a mature mangrove.

### Carbon as the master variable

Organic carbon was the strongest environmental signal in the dataset and the main constraint on community structure. Reference sediments held 2.5 and 5.5 times more organic carbon than altered and reforested sediments, and the increase in organic carbon and C/N with depth in the reference stand is consistent with long-term burial of root-derived, lignin-rich organic matter under anoxic conditions^2–4^. Arid mangroves such as Las Calderas often store less carbon than humid ones^33^, which makes the remaining stocks of intact stands particularly valuable. The taxa that tracked carbon point to the processes involved. Sediment Chloroflexota include anaerobic fermenters able to degrade complex and recalcitrant organic compounds^21^. Thermodesulfobacteriota comprise many sulfate reducers^22^, which typically dominate anaerobic carbon mineralisation in mangrove sediments^3,16,18^. Thermoproteota are archaea frequently associated with organic-rich anoxic layers. Together with the higher evenness of the reference community, this suggests a more complex, syntrophic anaerobic food web sustained by abundant organic substrates^17,19^.

The reforested stand had the lowest organic carbon of all stands, lower even than in the altered stand. Several mechanisms may contribute, including planting on previously cleared or eroded substrate, oxidation and export of legacy carbon after disturbance^4,38,39^, and limited root inputs from young trees. We also propose a priming hypothesis: labile inputs from young *R. mangle* litter and cyanobacterial necromass could stimulate the decomposition of older, more recalcitrant organic matter (positive priming)^40^, accelerating carbon loss rather than accumulation. This hypothesis is consistent with our data but requires direct tests, such as isotopic partitioning or incubations. The very low C/N ratios of reforested (1.3–2.6) and altered (2.5– 5.3) sediments are unusual for organic matter^41,42^. They suggest that, in these organic-carbon-poor sediments, total nitrogen includes a substantial inorganic fraction (e.g. clay-fixed ammonium), so these ratios should not be read as organic-matter stoichiometry alone.

### An open, alkaline sediment selects for phototrophs

The dominance of Cyanobacteriota in both disturbed stands, and their alignment with high pH in the ordinations, indicate that the loss of canopy turned the sediment surface into a habitat for benthic phototrophs. Without shade, light reaches the sediment and favours cyanobacterial mats^43,44^. Photosynthetic uptake of CO2 and bicarbonate by cyanobacteria raises porewater pH and can induce calcium carbonate precipitation^45,46^. This is consistent with the higher pH of the disturbed stands. The higher nitrogen of the altered stand may further favour fast-growing taxa, because nitrogen enrichment rapidly restructures mangrove sediment communities^42,47^. From a life-history perspective, nutrient-rich, disturbed conditions favour fast-growing (copiotrophic, r-selected) taxa, whereas slow-growing oligotrophs adapted to low substrate fluxes are outcompeted^48–50^. Many members of Chloroflexota, Thermodesulfobacteriota and Thermoproteota fit the latter strategy. The shift we observed may therefore reflect a transition from a slow, cooperative anaerobic community towards one dominated by fast-responding phototrophs. Acidobacteriota, which were most abundant in the altered stand, include oligotrophs able to use a wide range of carbon sources^51^ as well as lineages capable of dissimilatory sulfur metabolism^52^. They may therefore contribute to the degradation of residual organic matter in these sediments.

The persistence of high cyanobacterial read proportions below 30 cm warrants caution. Because the SILVA taxonomy places plastid 16S rRNA sequences within Cyanobacteriota^53^, all chloroplast and mitochondrial ASVs were removed before analysis, so this signal does not reflect algal or plant plastids. Buried mats, bioturbation by crabs and roots^54^, and relic extracellular DNA, which can persist in sediments and inflate the apparent abundance of inactive taxa^55^, may nonetheless contribute. Even under the most conservative interpretation, the shift from a carbon-processing to a phototroph-dominated profile distinguishes both disturbed stands from the reference forest.

### Sulfur and the anaerobic food web

Sulfur was nearly absent from the altered sediments, and sulfate-reducing Thermodesulfobacteriota were depleted in both disturbed stands. In intact mangroves, sulfate reduction coupled to the burial of iron sulfides retains sulfur in the sediment, and sulfur accumulates with depth as it did in our reference profile^3,18^. Physical disturbance reduces oxygen consumption and sulfate reduction rates in vegetated coastal sediments^56^. Lower organic-matter supply, oxidation of previously buried sulfides after canopy loss and possible losses of hydrogen sulfide could all deplete sediment sulfur. The relative importance of these pathways cannot be resolved with bulk elemental data, especially where cryptic sulfur cycling decouples sulfate reduction from net sulfide accumulation^57^. Porewater sulfate and sulfide profiles and acid-volatile sulfide measurements would clarify this question.

### Depth as a secondary structuring factor

Depth separated surface from subsurface communities, as reported for other soils and mangrove sediments in which oxygen, labile carbon and electron acceptors decline with depth^18,58,59^. In our data, the depth effect was weaker than the stand effect, and in the disturbed stands the vertical gradient was blurred by the persistence of Cyanobacteriota at depth. This compression of vertical structure is itself a symptom of degradation, because the redox stratification that organises mangrove sediment metabolism depends on continuous organic inputs from roots and litter^18,54^.

### Implications for restoration in the Caribbean

Our results have three practical implications. First, tree survival and canopy cover are insufficient indicators of restoration success. Sediment carbon and microbial community structure revealed that the reforested stand remained functionally equivalent to a degraded site. These below-ground indicators can be measured at modest cost and should be incorporated into national monitoring programmes^14,23^. Second, restoration in Las Calderas should address the conditions that limit organic-matter accumulation, notably tidal connectivity and the zonation of species, rather than relying on monospecific *R. mangle* plantings^9,13^. Third, the reference stand stores substantially more carbon than the disturbed stands and deserves strict protection as a carbon reservoir and as the benchmark against which restoration can be evaluated.

### Limitations

Aboveground recovery was assessed only qualitatively, from NDVI imagery (Supplementary Fig. S7); tree survival, canopy cover and basal area were not quantified, so the reforested stand is characterised here by its planting history and by its sediment and microbial properties. Our design pooled cores into composite samples. This reduces smallscale heterogeneity, but the reference stand was represented by a single composite per depth interval (pooled from three cores). Whole-stand contrasts therefore treat depth intervals as replicates, which are not independent, and depth-specific contrasts involving the reference stand lack replication^60^. PERMANOVA is sensitive to unequal dispersion in unbalanced designs^61^. Here the smaller group was the least dispersed, a configuration that tends to make the test conservative, but dispersion should be tested formally. Our inferences rely on relative abundances from rarefied amplicon data^62–64^. Quantitative PCR, compositionally aware differential-abundance methods^65^ and process measurements such as sulfate reduction rates or greenhouse-gas fluxes would strengthen functional interpretation. Finally, this is a single-time-point comparison. Replicated reference sites, a chronosequence of restoration ages and seasonal sampling are needed to determine whether, and how fast, reforested sediments converge on the reference state.

## Conclusion

In Las Calderas Bay, reforested mangrove sediments remained carbon-poor, alkaline and dominated by Cyanobacteriota. Their prokaryotic communities were indistinguishable from those of an altered mangrove and distinct from the carbon-rich, more diverse and anaerobic community of the reference forest. Sediment organic carbon, together with nitrogen, emerged as the main environmental driver of community structure, while depth acted as a secondary factor. Reforestation has therefore not yet been accompanied by belowground recovery. These findings provide the first microbial baseline for mangrove restoration in the Dominican Republic. They support incorporating sediment carbon and microbiome indicators into restoration monitoring, and designing interventions that restore the hydrological and sedimentary conditions needed for organic-matter burial.

## Acknowledgements

We thank the Ministerio de Medio Ambiente y Recursos Naturales of the Dominican Republic for granting access to genetic resources under contract VAPB-14918, the Instituto Especializado de Estudios Superiores Loyola for access to the elemental analyser, and Microbiome Insights (Vancouver, Canada) for amplicon sequencing. This paper was typeset with the bioRxiv Word template by @Chrelli: www.github.com/chrelli/bioRxiv-word-template.

## Author contributions

H.B.L.-C. conceived the study, designed the experiments, performed the elemental analyses, analysed the data in R and wrote the manuscript. J.A.C.-H. co-designed the study, analysed the data in R and supervised the work. A.N.A.-P., J.E.S.-J. and L.G.-E. carried out the fieldwork, sample processing and DNA extraction and contributed to data interpretation. All authors reviewed and approved the final manuscript.

## Permits and access to genetic resources

Sampling and access to genetic resources were carried out under the Contract for Access to Genetic Resources VAPB-14918 granted by the Ministerio de Medio Ambiente y Recursos Naturales of the Dominican Republic.

## Data and code availability

Raw sequence reads have been deposited in the NCBI Sequence Read Archive under BioProject PRJNA1538413 (BioSample accessions are listed in Supplementary Table S6). Sample metadata and sediment physicochemistry are provided in Supplementary Table S1. R scripts are available from the corresponding author upon request.

## Materials and Methods

### Study site

The study was conducted in Las Calderas Bay, Peravia province, on the arid south-western coast of the Dominican Republic (18.22° N, 70.51° W; Fig. 1). The site lies within the Las Dunas de Las Calderas Natural Monument (Decree 571-09; Law 202-04), a protected area of 17.48 km^2^. Its mangroves are dominated by *Rhizophora mangle*, with *Avicennia germinans, Laguncularia racemosa* and *Conocarpus erectus* in lower abundance. Although the area has been protected since 1996, reforestation did not begin until 2019 and has consisted of sporadic plantings of *R. mangle*^35^. Three stands were selected to represent different conservation states: a reference mangrove, a reforested stand and an altered stand affected by clearing and burning^34^. The conservation status of the reference stand was assessed by comparing Sentinel-2 Level-2A NDVI images from 2016 and 2025 in the Copernicus Browser (Supplementary Fig. S7).

### Sampling design

Sediment cores (1 m long) were collected on 17 January 2026 following the Coastal Blue Carbon protocols^66,67^. Eighteen cores were collected in each of the reforested and altered stands and three in the reference stand (39 cores). Each core was sectioned into four depth intervals: 0–15, 15–30, 30– 50 and 50–100 cm. To reduce small-scale spatial heterogeneity, a composite sampling strategy was used. In the reforested and altered stands, the 18 cores were randomly assigned to three independent groups of six. Sediment from the same depth interval within each group was homogenised, producing three independent composite samples per depth interval (B1–B3; n = 3). In the reference stand, sediment from each depth interval of the three cores was pooled into a single composite sample. In total, 28 composite samples were obtained: 12 reforested, 12 altered and 4 reference (Supplementary Table S1). Sampling areas were centred at 18.2224° N, 70.5119° W (reference), 18.2223° N, 70.5104° W (reforested) and 18.2191° N, 70.5090° W (altered). Sub-samples for DNA extraction were stored at −80 °C, and sub-samples for physicochemical analyses at −20 °C until processing.

### Physicochemical analyses

Sediments were dried at 60 °C for 72 h to constant weight. pH was measured in a 1:2.5 (w/v) sediment–water suspension following the FAO standard operating procedure. Salinity was calculated from the change in electrical conductivity after adding the sediment sample and expressed as NaCl (% w/w). Total dissolved solids (TDS) were also recorded. Total carbon (TC), hydrogen, nitrogen and sulfur contents were determined on 15–25 mg aliquots of dried, untreated whole sediment at IEESL with a Thermo Scientific FLASH 2000 organic elemental analyser in CHNS mode. The instrument was calibrated with 2–3 mg of sulfanilamide using the K-factor method^68^. Samples were not acidified, because acid pretreatment can artificially increase measured H and alter N and S concentrations in marine sediments^69^. Total inorganic carbon (TIC) was measured on a second aliquot after combustion at 500 °C to remove organic matter, and total organic carbon (TOC) was calculated as TC − TIC. Statistical analyses and C/N and C/S ratios used TOC. Sulfur values below the detection limit were recorded as zero; the C/S ratio is undefined for these samples and was set to zero in the analyses.

### DNA extraction, controls and sequencing

DNA was extracted from 0.25 g of each homogenised composite sample with the ZymoBIOMICS DNA Miniprep kit (D4300, Zymo Research, USA) following the manufacturer’s instructions. DNA concentration was measured with a NanoDrop spectrophotometer (Thermo Fisher Scientific, USA). The V4 region of the 16S rRNA gene was amplified with the universal primers 515F/806R^70^. The ZymoBIOMICS Microbial Community Standard (D6300)^71^ was processed as a positive control, and double-distilled water was processed as a negative control^72,73^. Amplification and paired-end sequencing were performed on the Illumina MiSeq platform at Microbiome Insights (Vancouver, Canada).

### Bioinformatic processing

Quality filtering, trimming, denoising, merging of paired reads, inference of amplicon sequence variants (ASVs) and chimera removal were carried out in R v4.6.1^74^ with DADA2 v1.40.0^75,76^. Forward and reverse reads were truncated at 260 and 200 bp, respectively. Reads with more than two expected errors (maxEE = 2) or with ambiguous bases were discarded. Taxonomy was assigned against the SILVA SSU database, release 138.2^53^. Contaminant ASVs were identified and removed with decontam v1.32.0^36^ using the negative control. Chloroplast and mitochondrial ASVs were removed. Samples were rarefied to 14,536 reads before diversity analyses^63,64^.

### Statistical analyses

Analyses were performed in R v4.6.1^74^ with microeco v2.3.0^77^, vegan v2.7-5^78^, phyloseq v1.56.0^79^ and microbiome v1.34.0^80^. Alpha diversity was estimated as observed ASV richness, Chao1^81^, the Shannon index^82^ and Pielou’s evenness^83^. Normality was checked with the Shapiro–Wilk test, and diversity indices were compared among stands, and among stand × depth groups, by one-way ANOVA followed by Tukey’s HSD. Community composition was compared using Bray–Curtis dissimilarities, visualised by NMDS and tested with pairwise PERMANOVA (adonis2, 999 permutations)^84^ between stands and between depth intervals, with Benjamini– Hochberg (BH) correction^85^. Differences in the relative abundance of phyla were tested with Wilcoxon rank-sum tests or Kruskal–Wallis tests followed by Dunn’s post hoc tests, as appropriate. Sediment variables were compared by ANOVA with Tukey’s HSD, by pairwise Wilcoxon rank-sum tests among stands (BH-corrected across all 24 tests) and by Kruskal–Wallis tests with Dunn’s comparisons among stand × depth groups.

For constrained ordinations, sediment variables were log_10_(x + 1)-transformed. dbRDA^86^ was performed on Bray–Curtis dissimilarities with N, TOC, H, S, C/N, C/S, pH and salinity as predictors. The significance of terms, added sequentially, and of constrained axes was assessed with 999 permutations. Phylum-level associations with sediment variables were examined by RDA and by Pearson correlations calculated within each stand. Statistical significance was set at α = 0.05.

