## Supplemental Information 1 for "Reforestation without Belowground Recovery: Sediment Microbiomes and Organic Carbon in Caribbean Mangroves"

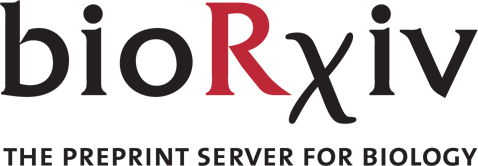

Supplementary Information

**for** “Reforestation without Belowground Recovery: Sediment Microbiomes and Organic Carbon in Caribbean Mangroves”

Hansel B. Lopez-Chevalier,^1,^* Andy N. Avelino-Perez,^2^ Juleidy E. Suriel-Jaime,^2^ Leslys Garcia-Encarnacion^2^ and Julian A. Cabrera-Hernandez^2^

^1^Instituto Especializado de Estudios Superiores Loyola (IEESL), San Cristóbal, Dominican Republic

^2^Facultad de Ciencias, Universidad Autónoma de Santo Domingo (UASD), Santo Domingo, Dominican Republic

### Contents

Supplementary Figure S1. Mock community and extraction blank

Supplementary Figure S2. Dominant phyla by stand and by stand × depth

Supplementary Figure S3. Alpha diversity by stand × depth

Supplementary Figure S4. Sediment physicochemistry by stand × depth

Supplementary Figure S5. Sediment physicochemistry along depth within stands

Supplementary Figure S6. Phylum–environment correlations within stands

Supplementary Figure S7. NDVI of the study area in 2016 and 2025

Supplementary Table S1. Sample metadata and sediment physicochemistry

Supplementary Table S2. Pairwise PERMANOVA

Supplementary Table S3. Pairwise Wilcoxon tests of sediment variables

Supplementary Table S4. dbRDA permutation tests

Supplementary Table S5. Mock community: expected vs observed composition

Supplementary Table S6. Sample codes, sequencing IDs and NCBI BioSample accessions

All tables are also provided in machine-readable form in Supplementary_Tables_S1-S6.xlsx. Sequence data: NCBI BioProject PRJNA1538413.

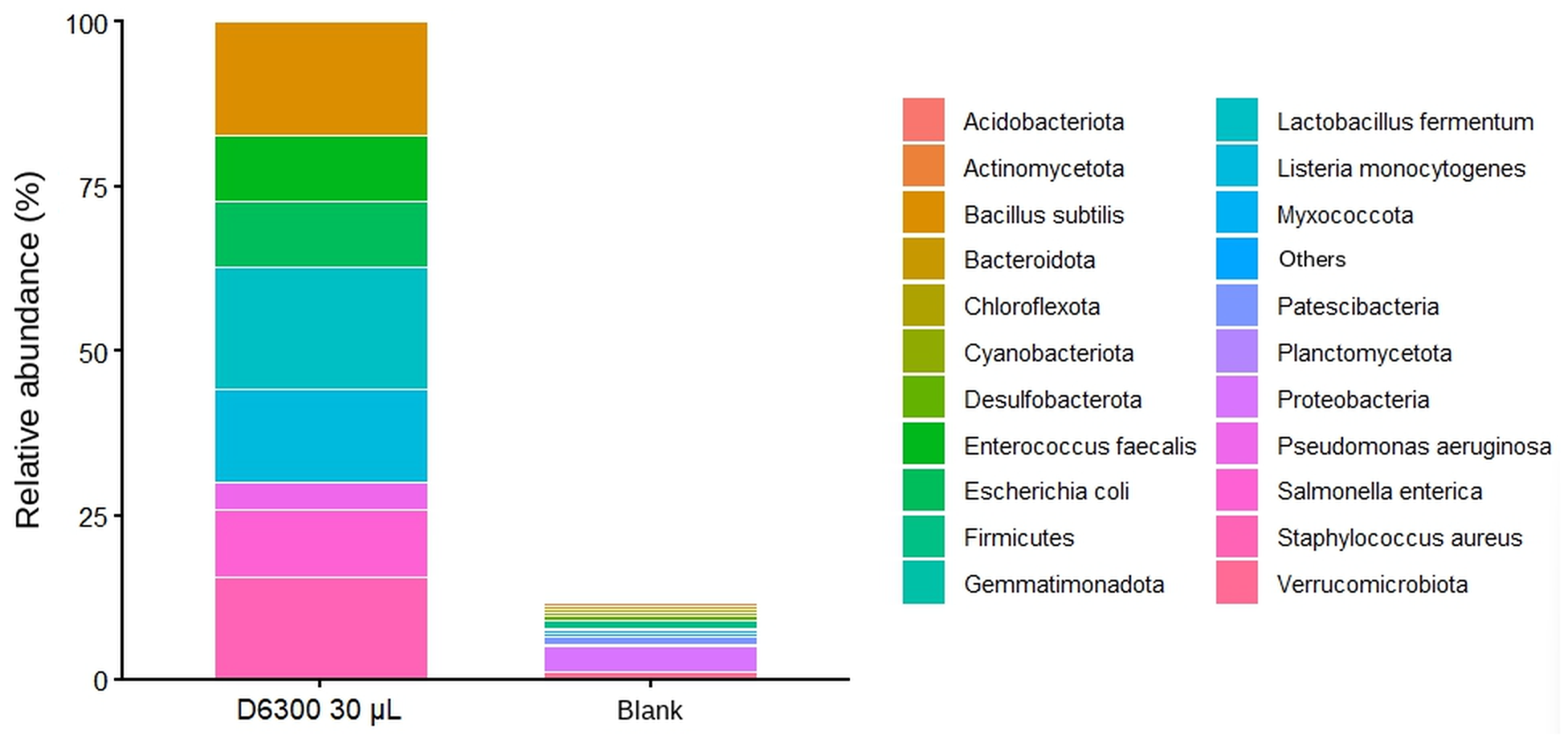

Supplementary Figure S1: Sequencing controls. Relative abundance of taxa in the ZymoBIOMICS Microbial Community Standard (D6300; 30 µL processed from extraction to sequencing) and in the negative control (double-distilled water). Mock members are shown at species level and environmental taxa at phylum level. All eight bacterial members of the mock were recovered in proportions matching the theoretical 16S rRNA gene composition (Supplementary Table S5). The negative control, plotted on the same scale as the mock library, contained only low-abundance environmental phyla and none of the mock taxa.

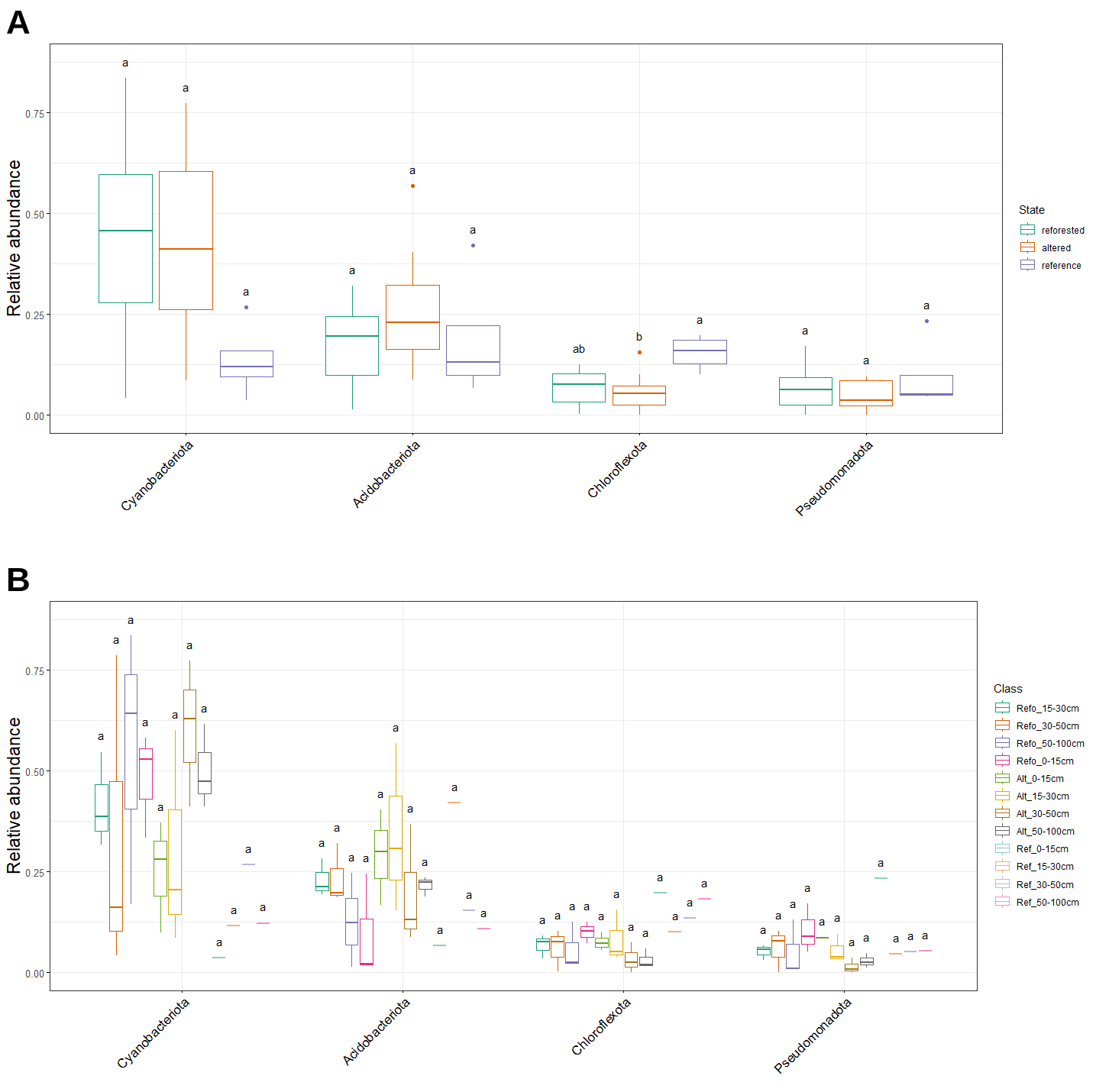

Supplementary Figure S2: Relative abundance of the four dominant phyla. **(A)** By stand and **(B)** by stand × depth group. Letters denote groups that differ significantly (ANOVA and Tukey’s HSD, *P* < 0.05). In (B), reference groups contain a single composite sample (pooled from three cores) and cannot be tested formally.

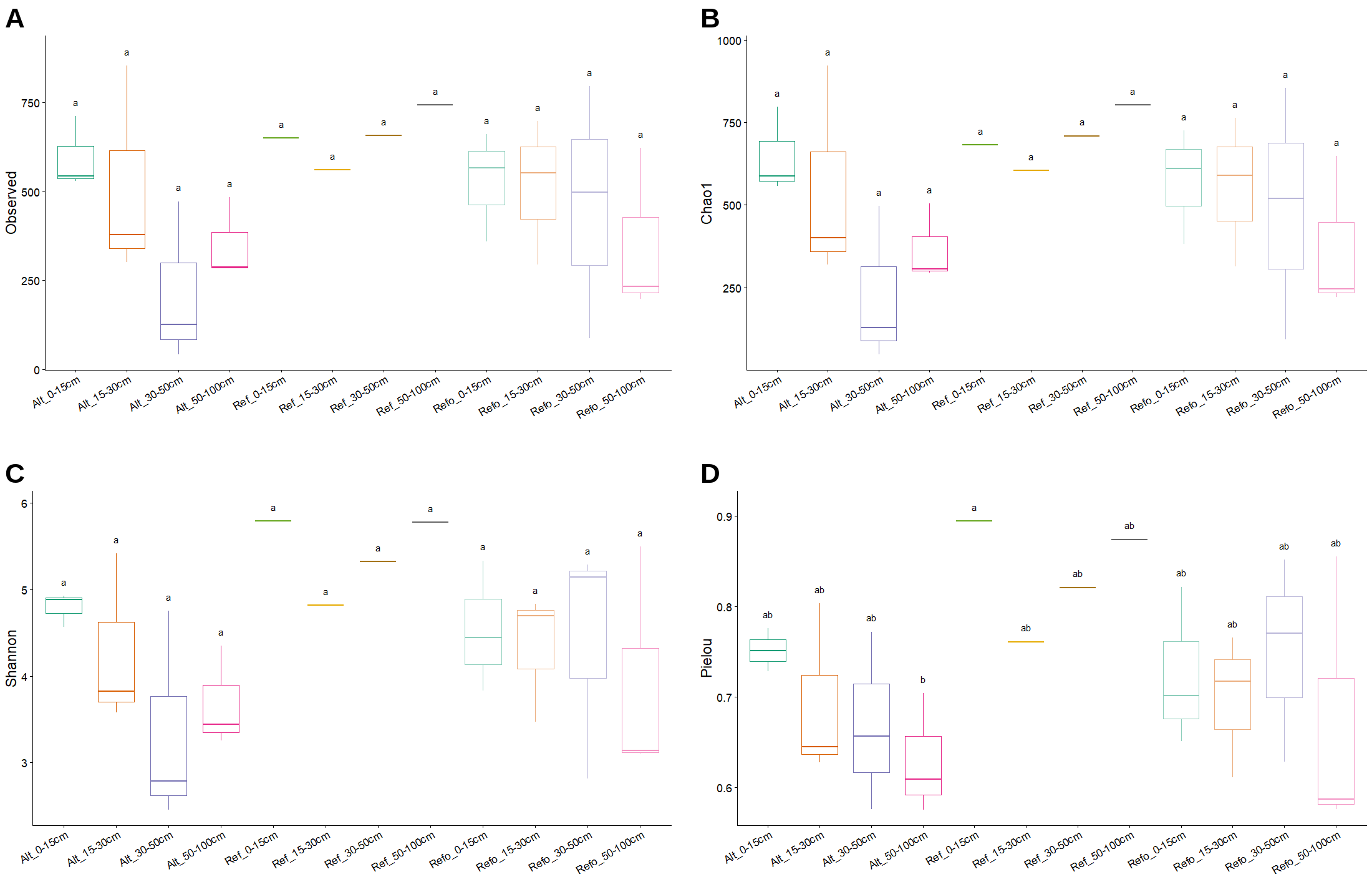

Supplementary Figure S3: Alpha diversity by stand × depth. **(A)** Observed ASV richness, **(B)** Chao1, **(C)** Shannon index and **(D)** Pielou’s evenness. Letters denote significant differences (ANOVA and Tukey’s HSD, *P* < 0.05). Reference groups (Ref_) contain one composite sample each (pooled from three cores); letters involving these groups should be interpreted with caution.

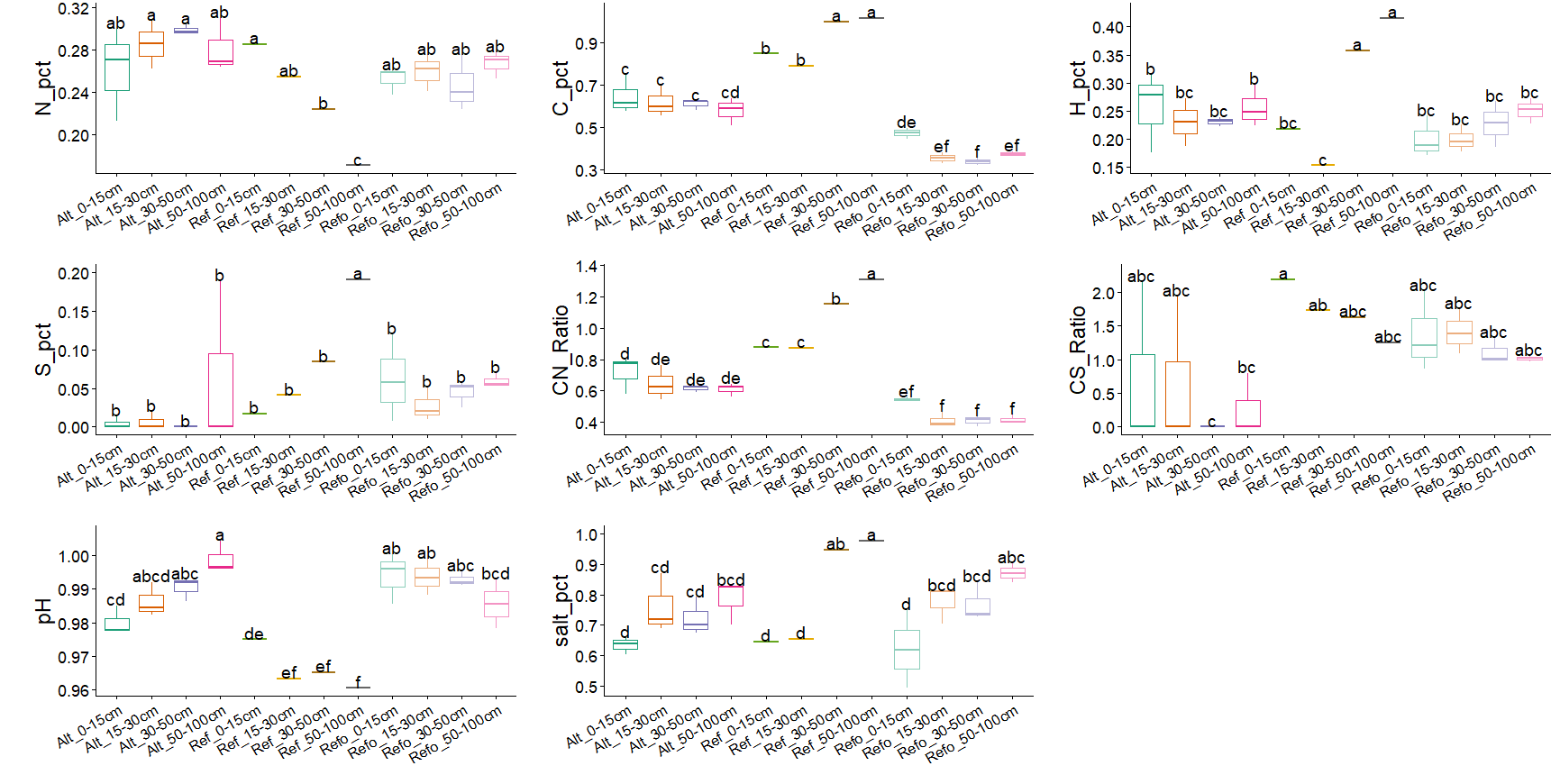

Supplementary Figure S4: Sediment physicochemistry by stand × depth. Variables were log_10_(x + 1)-transformed. Letters denote significant differences (ANOVA and Tukey’s HSD, *P* < 0.05). Reference groups contain one composite sample each. Kruskal–Wallis tests with Dunn’s post hoc comparisons detected no significant differences among stand × depth groups for any variable.

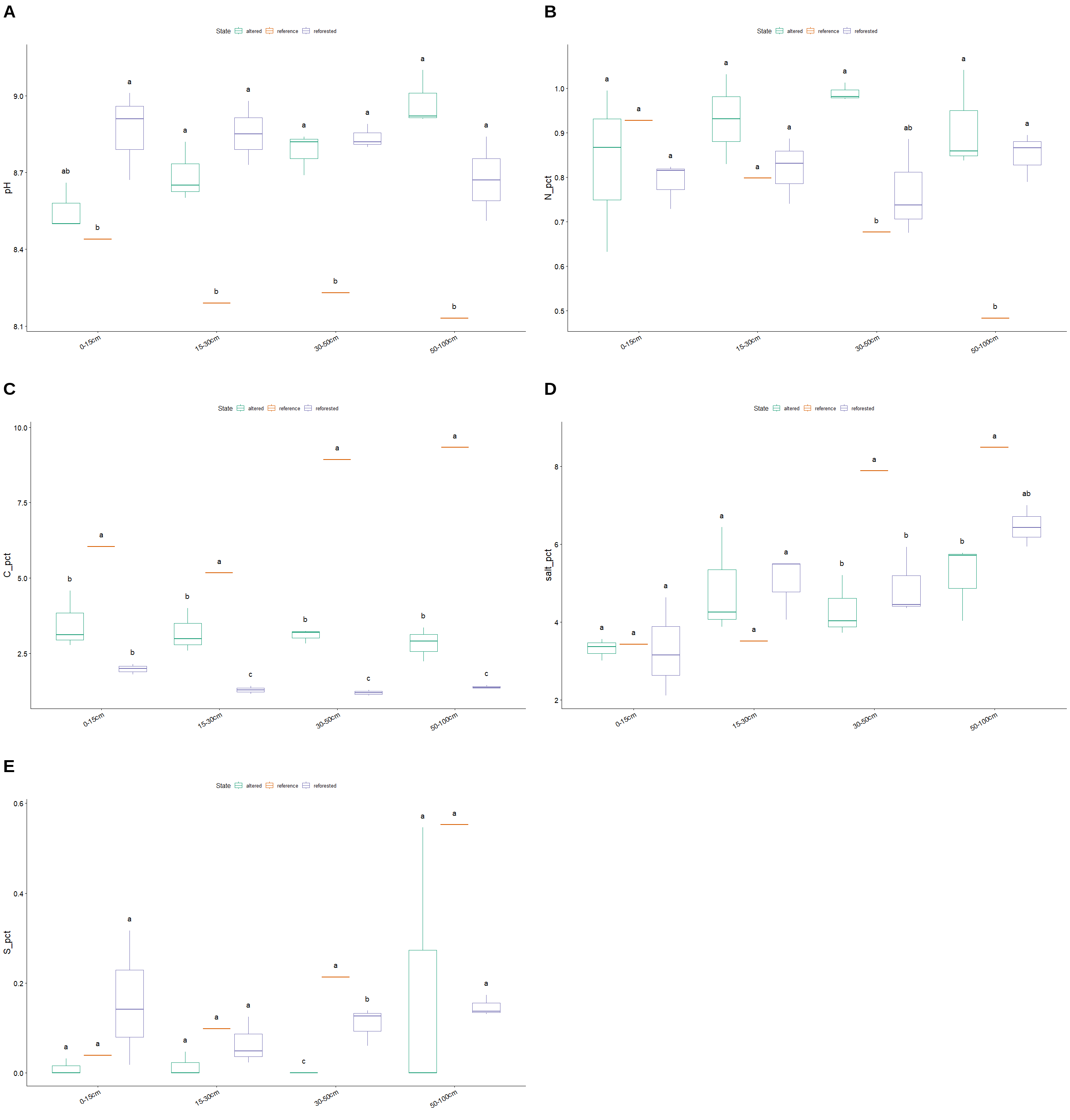

Supplementary Figure S5: Sediment physicochemistry along depth within each stand. **(A)** pH, **(B)** N (%), **(C)** C (%), **(D)** salinity (NaCl, %) and **(E)** S (%) at each depth interval in altered (n = 3 per depth), reference (one composite of three cores per depth) and reforested (n = 3 per depth) stands. Letters compare stands within each depth interval (ANOVA and Tukey’s HSD, *P* < 0.05).

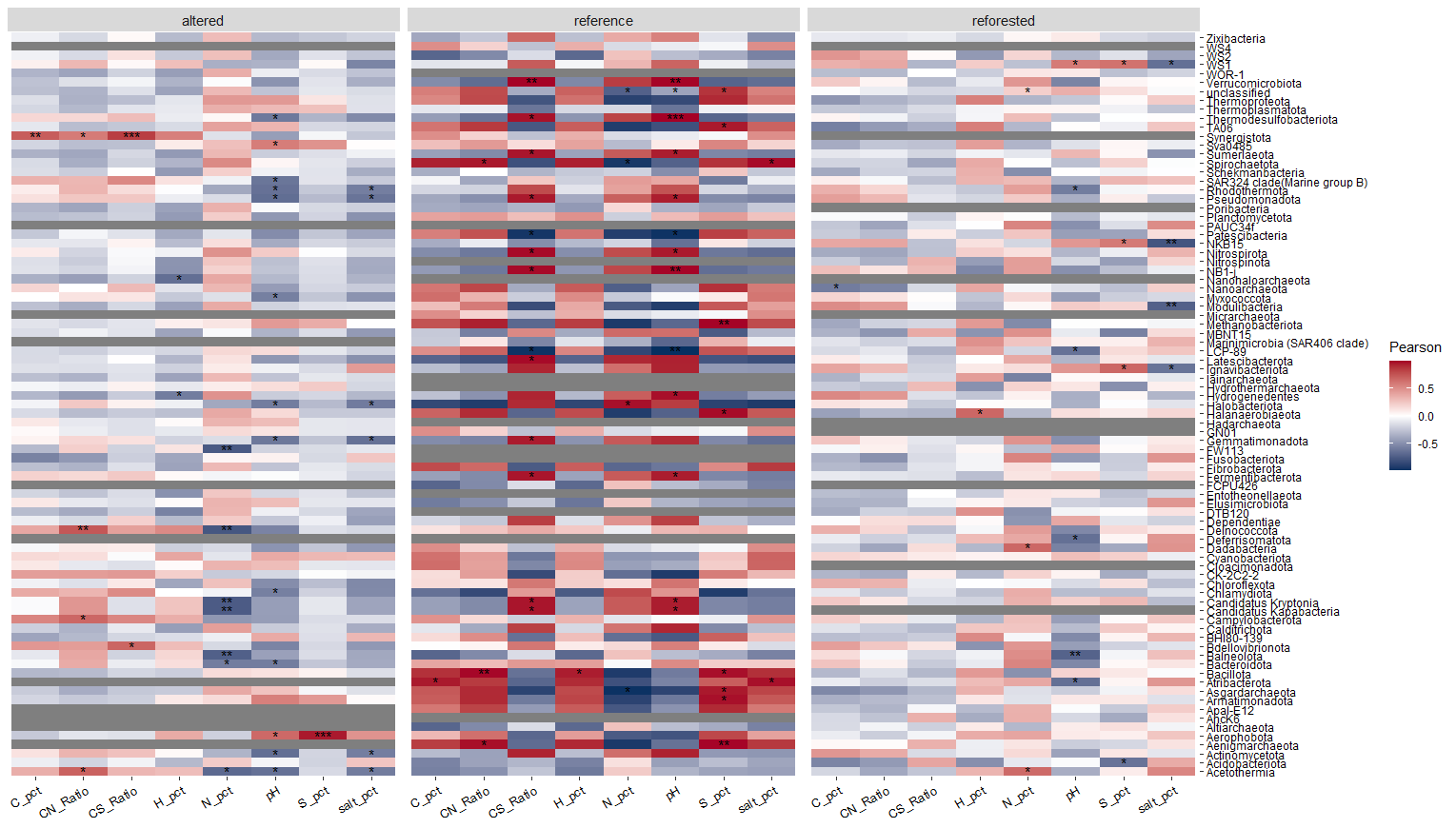

Supplementary Figure S6: Correlations between phyla and sediment variables within each stand. Heatmaps of Pearson correlation coefficients between the relative abundance of each phylum and sediment variables, calculated separately for altered (n = 12), reference (n = 4) and reforested (n = 12) samples. Grey cells, correlations not computable (taxon absent or invariant). * *P* < 0.05; ** *P* < 0.01; *** *P* < 0.001. Correlations for the reference stand are based on four samples and are shown for completeness only.

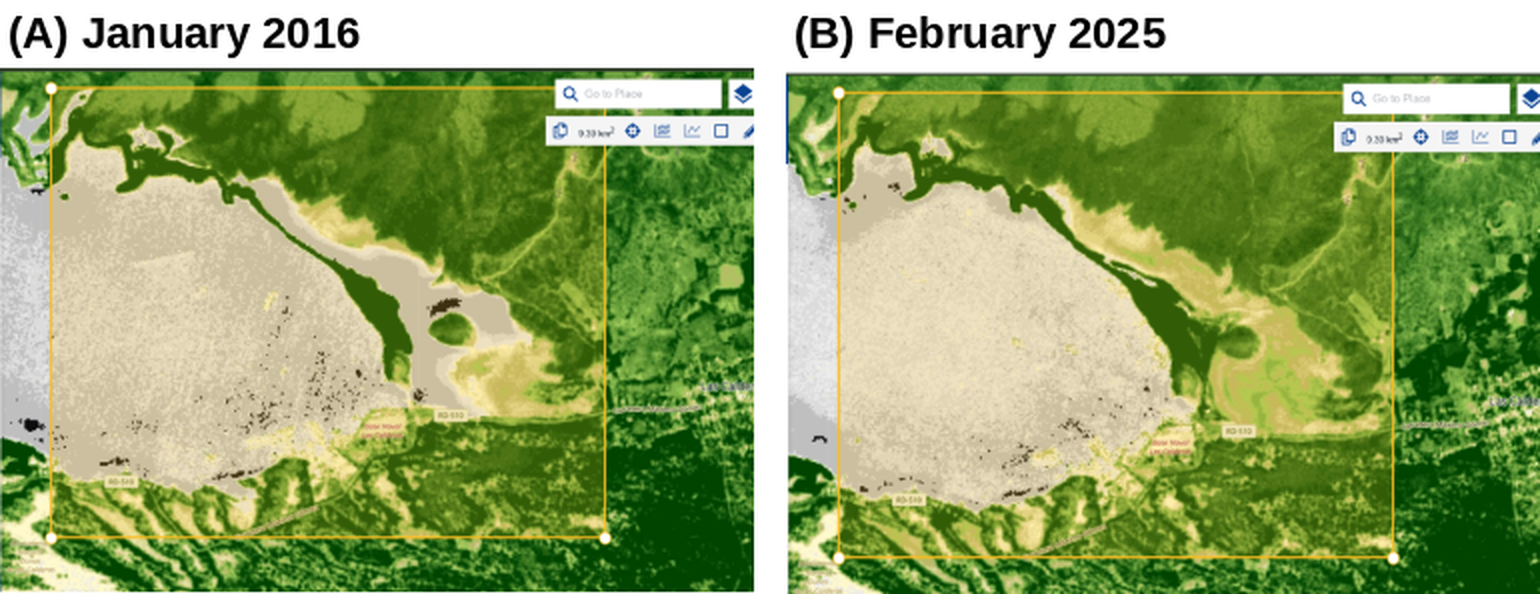

Supplementary Figure S7: Normalized Difference Vegetation Index (NDVI) of the study area. Sentinel-2 Level-2A NDVI, calculated as (B08 − B04)/(B08 + B04), on **(A)** 10 January 2016 and **(B)** February 2025, visualised in the Copernicus Browser (Change Detection through Time configuration; area of interest 9.39 km², yellow rectangle). Dark green indicates dense, photosynthetically active canopy and pale tones indicate bare sand, salt flats or water. The dense mangrove belt along the lagoon margin, where the reference stand is located, kept high NDVI values on both dates, supporting its use as a relatively undisturbed reference. Areas of low NDVI adjacent to this belt correspond to degraded or sparsely vegetated sediments; some of them show higher NDVI in 2025, consistent with incipient vegetation recovery.

Supplementary Table S1: Sample metadata and sediment physicochemistry.

| **Sample** | **Stand** | **Group** | **Depth (cm)** | **N** | **TOC** | **TIC** | **TC** | **H** | **S** | **C/N** | **C/S** | **pH** | **NaCl (%)** | **TDS** |
| --- | --- | --- | --- | --- | --- | --- | --- | --- | --- | --- | --- | --- | --- | --- |
| REFE-1 | Reference | — | 0–15 | 0.93 | 6.05 | 1.09 | 7.14 | 0.65 | 0.040 | 6.52 | 153.0 | 8.44 | 3.43 | 4.88 |
| REFE-2 | Reference | — | 15–30 | 0.80 | 5.17 | 1.40 | 6.57 | 0.42 | 0.099 | 6.47 | 52.4 | 8.19 | 3.51 | 4.44 |
| REFE-3 | Reference | — | 30–50 | 0.68 | 8.93 | 1.25 | 10.18 | 1.28 | 0.213 | 13.19 | 41.8 | 8.23 | 7.88 | 10.00 |
| REFE-4 | Reference | — | 50–100 | 0.48 | 9.33 | 2.05 | 11.38 | 1.60 | 0.552 | 19.31 | 16.9 | 8.13 | 8.48 | 10.00 |
| REFO-4 | Reforested | B1 | 0–15 | 0.82 | 1.99 | 0.62 | 2.61 | 0.74 | 0.317 | 2.42 | 6.3 | 9.01 | 2.11 | 2.99 |
| REFO-11 | Reforested | B1 | 15–30 | 0.74 | 1.42 | 0.23 | 1.65 | 0.68 | 0.125 | 1.92 | 11.4 | 8.98 | 5.49 | 7.80 |
| REFO-12 | Reforested | B1 | 30–50 | 0.74 | 1.28 | 0.31 | 1.59 | 0.70 | 0.139 | 1.74 | 9.3 | 8.89 | 4.36 | 6.05 |
| REFO-6 | Reforested | B1 | 50–100 | 0.79 | 1.44 | 0.17 | 1.61 | 0.69 | 0.174 | 1.82 | 8.3 | 8.67 | 5.94 | 8.43 |
| REFO-5 | Reforested | B2 | 0–15 | 0.81 | 2.14 | 0.62 | 2.76 | 0.54 | 0.141 | 2.63 | 15.2 | 8.67 | 3.15 | 2.48 |
| REFO-1 | Reforested | B2 | 15–30 | 0.83 | 1.15 | 0.23 | 1.38 | 0.57 | 0.049 | 1.38 | 23.3 | 8.85 | 4.06 | 5.76 |
| REFO-2 | Reforested | B2 | 30–50 | 0.89 | 1.19 | 0.40 | 1.59 | 0.54 | 0.060 | 1.35 | 19.9 | 8.80 | 5.93 | 8.69 |
| REFO-3 | Reforested | B2 | 50–100 | 0.89 | 1.36 | 0.23 | 1.59 | 0.79 | 0.131 | 1.52 | 10.4 | 8.51 | 7.00 | 9.94 |
| REFO-8 | Reforested | B3 | 0–15 | 0.73 | 1.79 | 0.47 | 2.26 | 0.49 | 0.017 | 2.46 | 102.6 | 8.91 | 4.63 | 6.58 |
| REFO-9 | Reforested | B3 | 15–30 | 0.89 | 1.28 | 0.17 | 1.45 | 0.51 | 0.023 | 1.44 | 54.6 | 8.73 | 5.50 | 7.78 |
| REFO-10 | Reforested | B3 | 30–50 | 0.67 | 1.09 | 0.31 | 1.40 | 0.85 | 0.126 | 1.62 | 8.7 | 8.82 | 4.45 | 6.53 |
| REFO-7 | Reforested | B3 | 50–100 | 0.87 | 1.33 | 0.28 | 1.61 | 0.87 | 0.138 | 1.53 | 9.7 | 8.84 | 6.42 | 9.07 |
| DETE-1 | Altered | B1 | 0–15 | 0.63 | 3.11 | 0.47 | 3.58 | 0.90 | 0.000 | 4.92 | n.d. | 8.50 | 3.01 | 4.18 |
| DETE-2 | Altered | B1 | 15–30 | 1.03 | 2.58 | 0.83 | 3.41 | 0.54 | 0.000 | 2.50 | n.d. | 8.60 | 3.88 | 4.86 |
| DETE-3 | Altered | B1 | 30–50 | 1.01 | 3.24 | 0.62 | 3.86 | 0.73 | 0.000 | 3.20 | n.d. | 8.69 | 5.20 | 7.46 |
| DETE-4 | Altered | B1 | 50–100 | 0.84 | 2.23 | 0.56 | 2.79 | 0.67 | 0.000 | 2.66 | n.d. | 8.91 | 5.71 | 8.09 |
| DETE-5 | Altered | B2 | 0–15 | 0.87 | 4.58 | 0.50 | 5.08 | 1.06 | 0.032 | 5.28 | 142.1 | 8.50 | 3.56 | 5.10 |
| DETE-9 | Altered | B2 | 15–30 | 0.83 | 4.00 | 0.92 | 4.92 | 0.87 | 0.047 | 4.83 | 85.2 | 8.65 | 4.25 | 6.05 |
| DETE-7 | Altered | B2 | 30–50 | 0.98 | 3.20 | 0.96 | 4.16 | 0.71 | 0.000 | 3.28 | n.d. | 8.82 | 4.03 | 5.81 |
| DETE-8 | Altered | B2 | 50–100 | 1.04 | 3.35 | 0.54 | 3.89 | 0.77 | 0.000 | 3.22 | n.d. | 8.92 | 4.03 | 5.76 |
| DETE-6 | Altered | B3 | 0–15 | 0.99 | 2.77 | 0.75 | 3.52 | 0.50 | 0.000 | 2.78 | n.d. | 8.66 | 3.37 | 4.84 |
| DETE-10 | Altered | B3 | 15–30 | 0.93 | 2.98 | 0.42 | 3.40 | 0.70 | 0.000 | 3.20 | n.d. | 8.82 | 6.43 | 9.05 |
| DETE-11 | Altered | B3 | 30–50 | 0.98 | 2.82 | 0.93 | 3.75 | 0.67 | 0.000 | 2.88 | n.d. | 8.84 | 3.72 | 5.28 |
| DETE-12 | Altered | B3 | 50–100 | 0.86 | 2.90 | 0.58 | 3.48 | 0.98 | 0.546 | 3.38 | 5.3 | 9.10 | 5.77 | 8.21 |

Samples were collected on 17 January 2026 in Las Calderas Bay (reference 18.2224° N, 70.5119° W; reforested 18.2223° N, 70.5104° W; altered 18.2191° N, 70.5090° W). Group: B1–B3 independent composite samples (each pooled from six cores) in reforested and altered stands; reference samples are composites of three cores per depth interval. Elemental contents are % dry weight. TC, total carbon measured on untreated sediment; TIC, inorganic carbon measured on the residue after combustion at 500 °C; TOC = TC − TIC. Statistical analyses and C/N and C/S ratios used TOC. n.d., C/S not defined because S was below the detection limit (recorded as 0 in the original dataset). Note: in the original spreadsheet the columns for S and H were labelled in reverse; values are reported here with corrected labels, consistent with the C/S ratio and with the analyses in R.

Supplementary Table S2: Pairwise PERMANOVA on Bray–Curtis dissimilarities (999 permutations).

| **Factor** | **Comparison** | **Pseudo-F** | **R²** | **P** | **Adjusted P** | **Sig.** |
| --- | --- | --- | --- | --- | --- | --- |
| Stand | Reforested vs altered | 1.023 | 0.044 | 0.252 | 0.252 |  |
| Stand | Reforested vs reference | 1.383 | 0.090 | 0.001 | 0.0015 | ** |
| Stand | Altered vs reference | 1.430 | 0.093 | 0.001 | 0.0015 | ** |
| Depth | 15–30 vs 30–50 cm | 1.019 | 0.078 | 0.344 | 0.413 |  |
| Depth | 15–30 vs 50–100 cm | 1.062 | 0.081 | 0.096 | 0.144 |  |
| Depth | 0–15 vs 15–30 cm | 1.097 | 0.084 | 0.037 | 0.074 |  |
| Depth | 30–50 vs 50–100 cm | 0.936 | 0.072 | 0.841 | 0.841 |  |
| Depth | 0–15 vs 30–50 cm | 1.204 | 0.091 | 0.007 | 0.021 | * |
| Depth | 0–15 vs 50–100 cm | 1.208 | 0.091 | 0.002 | 0.012 | * |

P values were adjusted with the Benjamini–Hochberg method within each factor. * adjusted *P* < 0.05; ** adjusted *P* < 0.01.

Supplementary Table S3: Pairwise Wilcoxon rank-sum tests of sediment variables among stands.

| **Variable** | **Comparison** | **P** | **Adjusted P** | **Sig.** | **Higher in** |
| --- | --- | --- | --- | --- | --- |
| N (%) | Refo–Alt | 0.010 | 0.020 | * | Alt |
| N (%) | Refo–Ref | 0.521 | 0.658 | n.s. | Refo |
| N (%) | Alt–Ref | 0.042 | 0.072 | n.s. | Alt |
| TOC (%) | Refo–Alt | 7.40×10^-7^ | 1.78×10^-5^ | *** | Alt |
| TOC (%) | Refo–Ref | 0.001 | 0.004 | ** | Ref |
| TOC (%) | Alt–Ref | 0.001 | 0.004 | ** | Ref |
| H (%) | Refo–Alt | 0.219 | 0.309 | n.s. | Alt |
| H (%) | Refo–Ref | 0.599 | 0.684 | n.s. | Ref |
| H (%) | Alt–Ref | 0.862 | 0.862 | n.s. | Ref |
| S (%) | Refo–Alt | 9.97×10^-4^ | 0.004 | ** | Refo |
| S (%) | Refo–Ref | 0.599 | 0.684 | n.s. | Ref |
| S (%) | Alt–Ref | 0.009 | 0.020 | * | Ref |
| C/N | Refo–Alt | 1.48×10^-6^ | 1.78×10^-5^ | *** | Alt |
| C/N | Refo–Ref | 0.001 | 0.004 | ** | Ref |
| C/N | Alt–Ref | 0.001 | 0.004 | ** | Ref |
| C/S | Refo–Alt | 0.004 | 0.010 | * | Refo |
| C/S | Refo–Ref | 0.058 | 0.093 | n.s. | Ref |
| C/S | Alt–Ref | 0.019 | 0.036 | * | Ref |
| pH | Refo–Alt | 0.340 | 0.453 | n.s. | Refo |
| pH | Refo–Ref | 0.004 | 0.010 | * | Refo |
| pH | Alt–Ref | 0.004 | 0.010 | * | Alt |
| NaCl (%) | Refo–Alt | 0.194 | 0.291 | n.s. | Refo |
| NaCl (%) | Refo–Ref | 0.684 | 0.713 | n.s. | Ref |
| NaCl (%) | Alt–Ref | 0.671 | 0.713 | n.s. | Ref |

P values were adjusted with the Benjamini–Hochberg method across all 24 tests. “Higher in” indicates the stand with the larger median. Refo, reforested; Alt, altered; Ref, reference.

Supplementary Table S4: Permutation tests for the dbRDA model.

| **Test** | **Term / axis** | **df** | **Sum of squares** | **F** | **P** |
| --- | --- | --- | --- | --- | --- |
| Term | N | 1 | 0.5198 | 1.094 | 0.047* |
| Term | TOC | 1 | 0.5684 | 1.197 | 0.005** |
| Term | H | 1 | 0.4806 | 1.012 | 0.397 |
| Term | S | 1 | 0.4770 | 1.004 | 0.455 |
| Term | C/N | 1 | 0.4078 | 0.858 | 1.000 |
| Term | C/S | 1 | 0.4521 | 0.952 | 0.838 |
| Term | pH | 1 | 0.4740 | 0.998 | 0.487 |
| Term | NaCl | 1 | 0.4732 | 0.996 | 0.458 |
| Term | Residual | 19 | 9.0255 | — | — |
| Axis | dbRDA1 | 1 | 0.6813 | 1.438 | 0.008** |
| Axis | dbRDA2 | 1 | 0.5380 | 1.135 | 0.779 |
| Axis | dbRDA3 | 1 | 0.4758 | 1.004 | 1.000 |
| Axis | Residual | 19 | 9.0045 | — | — |

Model: Bray–Curtis ~ N + TOC + H + S + C/N + C/S + pH + NaCl (log_10_(x + 1)-transformed predictors; 999 permutations). Terms were added sequentially (type I tests), so results depend on the order of entry. Term and axis tests come from two separate runs of the model (residual SS 9.0255 and 9.0045, respectively); in the second run N was marginal (*P* = 0.055).

Supplementary Table S5: Composition of the ZymoBIOMICS D6300 mock community.

| **Taxon** | **Expected (%)** | **Observed (%)** | **Difference (pp)** |
| --- | --- | --- | --- |
| *Pseudomonas aeruginosa* | 4.2 | 4.6 | +0.4 |
| *Escherichia coli* | 10.1 | 9.7 | -0.4 |
| *Salmonella enterica* | 10.4 | 10.8 | +0.4 |
| *Limosilactobacillus fermentum* | 18.4 | 17.9 | -0.5 |
| *Enterococcus faecalis* | 9.9 | 10.3 | +0.4 |
| *Staphylococcus aureus* | 15.5 | 14.8 | -0.7 |
| *Listeria monocytogenes* | 14.1 | 14.6 | +0.5 |
| *Bacillus subtilis* | 17.4 | 17.3 | -0.1 |

Expected values are the theoretical 16S rRNA gene composition provided by the manufacturer. Observed values were obtained from the DADA2/SILVA 138.2 output of the mock library.

Supplementary Table S6: Sample codes, sequencing IDs and NCBI accessions.

| **Lab code** | **Sequencing ID** | **Stand** | **Depth (cm)** | **BioSample** |
| --- | --- | --- | --- | --- |
| REFE-1 | S00UT-0025 | Reference | 0–15 | SAMN63852817 |
| REFE-2 | S00UT-0026 | Reference | 15–30 | SAMN63852818 |
| REFE-3 | S00UT-0027 | Reference | 30–50 | SAMN63852819 |
| REFE-4 | S00UT-0028 | Reference | 50–100 | SAMN63852820 |
| REFO-4 | S00UT-0004 | Reforested | 0–15 | SAMN63852793 |
| REFO-11 | S00UT-0011 | Reforested | 15–30 | SAMN63852794 |
| REFO-12 | S00UT-0012 | Reforested | 30–50 | SAMN63852795 |
| REFO-6 | S00UT-0006 | Reforested | 50–100 | SAMN63852796 |
| REFO-5 | S00UT-0005 | Reforested | 0–15 | SAMN63852797 |
| REFO-1 | S00UT-0001 | Reforested | 15–30 | SAMN63852798 |
| REFO-2 | S00UT-0002 | Reforested | 30–50 | SAMN63852799 |
| REFO-3 | S00UT-0003 | Reforested | 50–100 | SAMN63852800 |
| REFO-8 | S00UT-0008 | Reforested | 0–15 | SAMN63852801 |
| REFO-9 | S00UT-0009 | Reforested | 15–30 | SAMN63852802 |
| REFO-10 | S00UT-0010 | Reforested | 30–50 | SAMN63852803 |
| REFO-7 | S00UT-0007 | Reforested | 50–100 | SAMN63852804 |
| DETE-1 | S00UT-0013 | Altered | 0–15 | SAMN63852805 |
| DETE-2 | S00UT-0014 | Altered | 15–30 | SAMN63852806 |
| DETE-3 | S00UT-0015 | Altered | 30–50 | SAMN63852807 |
| DETE-4 | S00UT-0016 | Altered | 50–100 | SAMN63852808 |
| DETE-5 | S00UT-0017 | Altered | 0–15 | SAMN63852809 |
| DETE-9 | S00UT-0021 | Altered | 15–30 | SAMN63852810 |
| DETE-7 | S00UT-0019 | Altered | 30–50 | SAMN63852811 |
| DETE-8 | S00UT-0020 | Altered | 50–100 | SAMN63852812 |
| DETE-6 | S00UT-0018 | Altered | 0–15 | SAMN63852813 |
| DETE-10 | S00UT-0022 | Altered | 15–30 | SAMN63852814 |
| DETE-11 | S00UT-0023 | Altered | 30–50 | SAMN63852815 |
| DETE-12 | S00UT-0024 | Altered | 50–100 | SAMN63852816 |
| NEG-CTRL | S00UT-0029 | Negative control | — | SAMN63852821 |
| MOCK-D6300 | S001Z-0016 | Positive control | — | SAMN63852822 |

All data are available under NCBI BioProject PRJNA1538413. Lab code, sample identifier used in this study; Sequencing ID, Microbiome Insights sample identifier (FASTQ prefix).
